# Celldega: Integrated Toolkit for Visualization and Analysis of Spatial Data

**DOI:** 10.64898/2026.08.13.744672

**Authors:** Nicolas Fernandez, Jaspreet Ishar, Huan Wang, Amin Ben Saad, Michal Lipinski, Samouil L. Farhi

## Abstract

Spatial-transcriptomics integrates high-dimensional single-cell data with microscopy to reveal cellular states, communication, and tissue organization. Analyzing this data requires a combination of multi-modal data processing, high-dimensional data analysis, spatial analysis, and integrated visualization. However, computational analysis is increasingly becoming a bottleneck as approaches mature and dataset sizes increase. Additionally, visualization can be challenging as open-source visualization tools struggle to scale to large datasets (exceeding 1 billion transcripts), and commercial visualization tools are costly, closed source, and inflexible. We present Celldega, an open-source Python and JavaScript library for scalable, interactive visualization and analysis of spatial-omics data. Celldega integrates custom analyses, performs neighborhood analysis, implements an efficient visualization-specific file format, and enables interactive exploration in notebooks and web galleries. We demonstrate Celldega across multiple technologies, tissues, and datasets, including 3D reconstructions of the developing whole mouse head comprising over four million cells. Finally, we demonstrate how Celldega can be utilized throughout the entire lifecycle of spatial data analysis, from quality control to building a public shareable gallery.

## Intro

Spatial transcriptomics (ST) measures gene expression directly within intact tissue, preserving the native tissue context and microenvironment (1). This context enables exploration of cell morphology, cell-cell communication, cellular neighborhoods, tissue organization, and spatial gradients that are lost in dissociative single-cell transcriptomics. As experimental methods continue to advance, the data generated is covering increasingly larger areas, at finer resolution, and with higher molecular plex, with full-transcriptome profiling at subcellular resolution becoming the standard in the field. Fueled by these advances, both the number and scale of spatial studies are growing across research and into clinical settings.

As rapidly as the experimental methods have developed, the analysis community has responded with an even larger proliferation of methods, solving tasks from the lowest level processing of raw data to the most abstract AI agents powered by foundational spatial models (2). The richness of the data contributes to this excitement. At its core, spatial omics couples morphological and highly plexed molecular data, combining the demands and the opportunities of image processing with the high-dimensional matrix analysis of single-cell transcriptomics. A full spatial analysis frequently requires both imaging specific steps (e.g. cell identification and segmentation, feature extraction, region of interest selection) and expression analysis (e.g. cell type identification and clustering; differential expression analysis). Since the data is coupled, these analyses are better performed in tandem. We have seen already, for example, that segmentation is improved if expression information is factored in (3,4) and that ROIs can be selected based on gene expression (5). Emergent information is also available by joint analysis of these domains: identification of cell neighborhoods and microdomains (6); spatial co-expression analysis; and discovery of long range spatially varying genes (7). The field is actively exploring each of these avenues, bringing new computational methods regularly. Thus, despite several ongoing efforts (8–12) the process has not been standardized.

The biologist leveraging spatial biology is thus faced with a continually growing set of data formats and integrative data analysis challenges requiring multiple bespoke pieces of code which must work across imaging and matrix domains. The response is to work in teams, usually consisting of a combination of a biological area expert, a pathologist, and a computational expert to extract the most meaning from the data. Unfortunately, the available infrastructure to support team-based, flexible, multi-domain analyses across the imaging and transcriptomics domains is still lacking.

Existing open-source visualization tools often emphasize a single perspective – spatial- visualization (5,13), data-visualization (14–16), or customizable dashboards (17), with the latter often difficult to configure and less well suited to interactive notebook-based analysis. Open-source visualization tools can also struggle to render transcript-level data for large datasets (> one billion transcripts) (5,13,16,17), and often maintain responsiveness by limiting transcript visualization to selected genes, subsamples, or aggregates (18,19). Commercial visualization platforms from technology providers and cloud-computing platforms can offer additional scalability but are generally more restrictive and may also be desktop-only, closed-source, vendor-specific, or tied to hosted systems.

Finally, most spatial analysis workflows remain focused on single-cell-level analysis and individual samples, lacking explicit data structures to support the analysis of higher-order or hierarchical biological entities such as neighborhoods, tissue domains, gene modules, and entire cohorts. These challenges stem from fragmented tool development: data formats, analysis workflows, and visualizations are often developed in isolation, preventing seamless integration. As technologies advance, datasets grow, and large cohort-scale studies become common, these challenges will only increase.

To address these challenges, we developed Celldega – an integrated, scalable, and extensible framework that enables users to: visually explore and annotate spatial data throughout the research lifecycle, analyze higher-order biological structures (e.g., neighborhoods), and share results through public galleries and/or notebooks to promote open and reusable science. Celldega is named after a bodega, a small shop with all the essentials that is part of the fabric of a neighborhood. In the same spirit, Celldega brings together the essential views needed to explore spatial biology—from sub-cellular to tissue- level neighborhoods—pairing data and information visualization with the supporting data structures and methods for spatial analysis.

## Results

### Celldega’s design facilitates scalable spatial data visualization in notebooks and web galleries

Our primary goal in designing Celldega was to connect analysis and visualization of spatial data. We reasoned that if this integration could be sufficiently scalable, responsive, and customizable, a user would be empowered to take on new datasets and new analyses rapidly without the hurdles of existing tools. Therefore, the core of Celldega’s architecture revolves around a series of visualization tools and supporting data structures and methods (Fig. 1A). Built in visualizations span “physical”-space (e.g. images, spatially resolved data points); “data”-space (e.g. heatmaps, scatter plots); and “information”-space (e.g. enrichment analysis) (Fig. 1A(i)). Celldega assumes that most analysis carried out by the user will be performed with other software packages, and the tool is configured to be extensible to allow direct incorporation of analysis approaches by the user (e.g., using AnnData). Similarly, the focus is on using pre-existing data structures and formats where possible, with new data structures generated only as necessary to bridge gaps in performance (Fig. 1A(ii)). In particular, because spatial analysis often requires reasoning about structures above the single-cell level—neighborhoods, tissue domains, manually annotated regions—we developed methods for constructing neighborhoods, calculating neighborhood-level feature spaces, and visualizing neighborhoods, together with a class for storing neighborhoods and their feature spaces (Fig. 1A(iii)). Neighborhoods are designed to be maximally flexible and extendible to other tools. They can be constructed by manual annotation, tessellation of the tissue area, alpha shape (20), or gradient methods; they can be visualized from external sources such as third-party domain algorithms; and characterized by multi-omic feature spaces such as neighborhood-by-population matrices. To enable interactive analysis and visualization, Celldega is built on modern JavaScript, with ES6 modules rather than frameworks such as React, and Jupyter widgets, which enable simple deployment to either stand-alone galleries or notebook-based analyses. In notebooks, the Jupyter Widgets framework enables two-way communication between front-end visualizations and the Python kernel, as well as between widgets (Fig. 1B). This allows users to annotate data directly in visualizations while preserving the annotation state in Python. Widget linkage also allows Celldega views to communicate with each other or with third-party widgets such as jupyter-scatter (15), enabling spatial views, gene expression plots, and embedding visualizations to operate as coordinated parts of the same analysis (Fig. 1B). Finally, this infrastructure also allows for the extension to new data visualization widgets as demanded by the analysis.

**Fig. 1.**
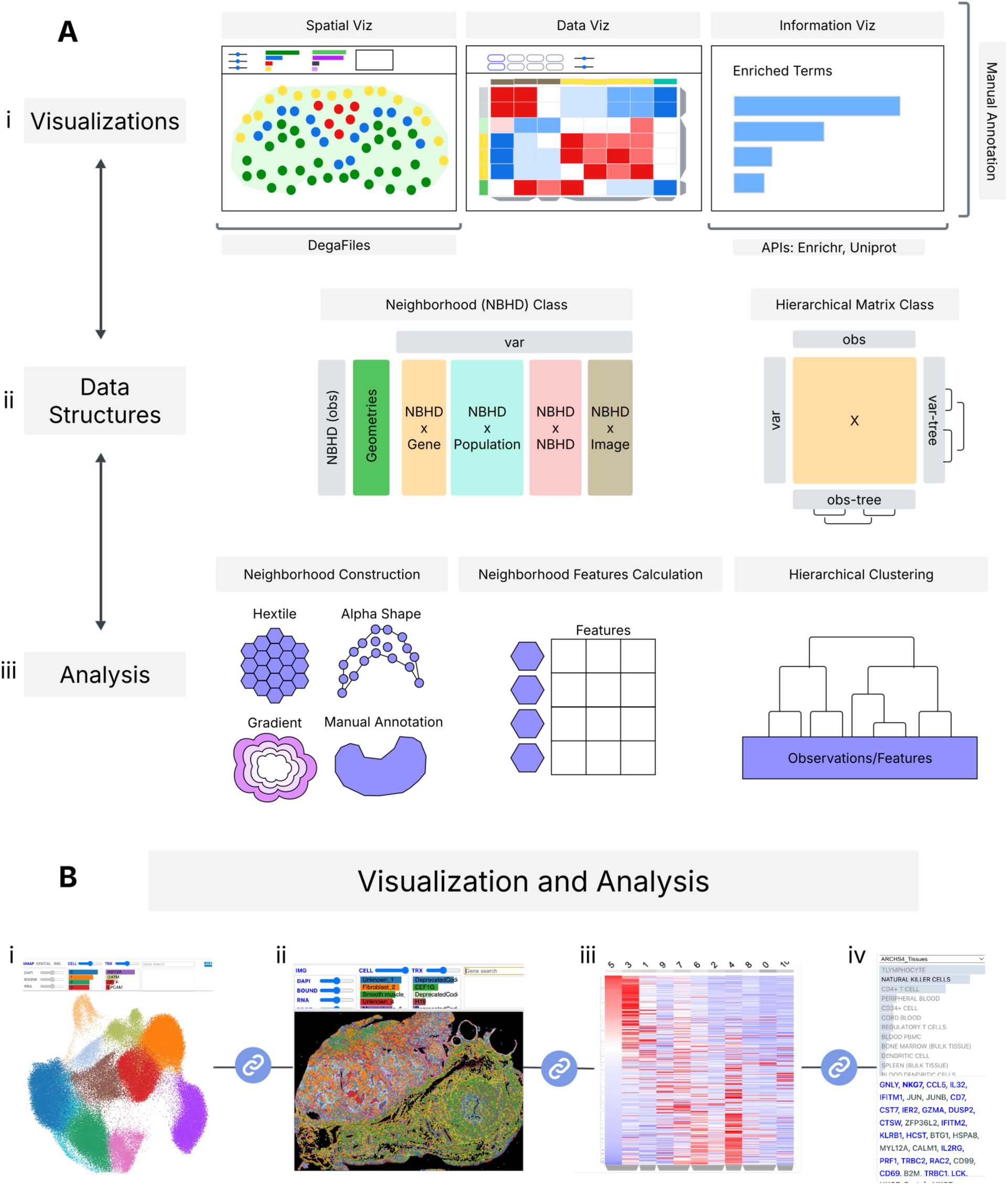
Celldega’s architecture couples interactive visualization with supporting data structures and analysis methods for spatial data. (A) Overview of Celldega’s three layers. (A(i)) Visualizations span physical space, data space, and information space; manual annotations can be performed across these views. DegaFiles enable spatial visualizations. (A(ii)) Data structures: the neighborhood class (NBHD) stores geometries and feature spaces. The hierarchical matrix class (Matrix) stores hierarchical biclustering results from observations (entities) and variables (features). (A(iii)) Analysis methods construct neighborhoods, calculate neighborhood-level feature spaces, and perform hierarchical clustering of observations and variables. (B) Celldega’s widgets can be linked to one another to support coordinated spatial data analysis. The Widget framework enables linking separate widgets: (B(i)) UMAP Landscape, (B(ii)) Spatial Landscape, (B(iii)) Clustergram, and (B(iv)) Enrich.

Underpinning our spatial visualizations are DegaFiles, a visualization-specific file architecture designed to minimize the computational overhead of sharing and visualizing large spatial transcriptomics datasets (Fig. 1A(i), left). DegaFiles organize the core components needed for interactive visualization, including multichannel images, segmented cell polygons, cell-by-gene expression data, cell and gene metadata, and transcript coordinates. Images are stored in the cloud-friendly WebP format, enabling efficient rendering of multiple layered channels with adjustable opacity. Inspired by developments in geospatial data systems, including GeoParquet, we developed a cloud-optimized Parquet tiling strategy for storing vector geometries such as transcripts and cell boundaries. Together, this file architecture enables a fully client-side, serverless model for interactive visualization of large spatial datasets. Celldega generates DegaFiles from raw instrument data from many of the leading commercial ST technologies including: Xenium, CosMx, MERSCOPE, Visium HD, StrataMap, and Atera (Fig. 2; Supp. Fig. 1). To future proof Celldega we have included documentation on formatting a new reader and plan to support SpatialData as an intermediate format to streamline the support of additional technologies—even to homebrew methods, such as a 100 μm thick MERFISH dataset (Supp. Fig. 1D).

**Fig. 2.**
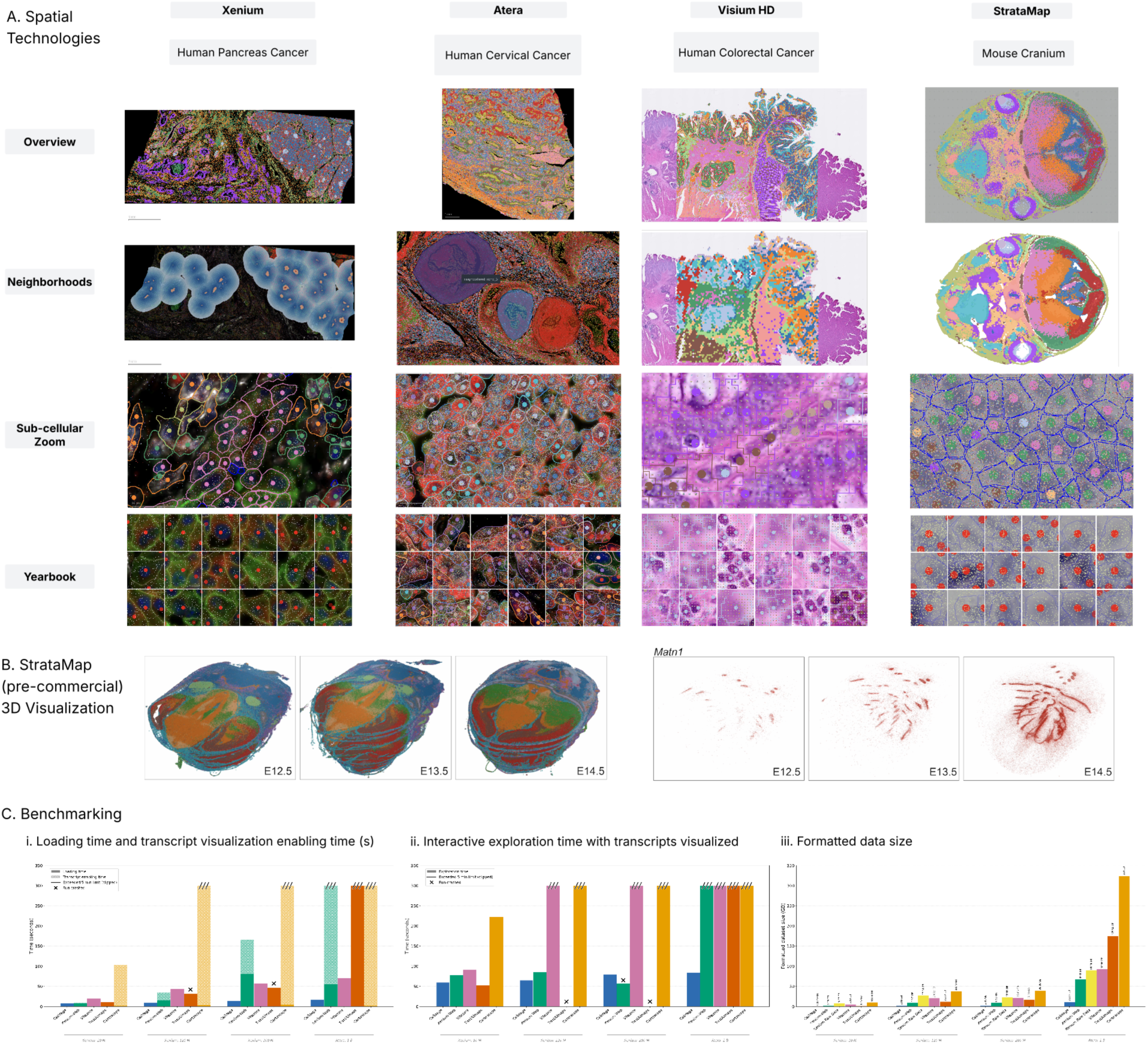
Celldega supports scalable visualization of large spatial transcriptomics datasets across imaging- and sequencing-based technologies. (A) Multi-scale visualization of four datasets spanning distinct spatial platforms: Xenium (human pancreatic cancer), Atera (human cervical cancer), Visium HD (human colorectal cancer), and StrataMap (pre-commercial) (developing whole mouse head, E14.5). Each column shows, from top to bottom: a whole-sample overview; neighborhood visualizations— gradient (Xenium), manual annotation (Atera), hextile (Visium HD), and alpha shape (StrataMap); a sub-cellular view resolving individual cell boundaries and transcripts (a small jitter, e.g., 1 µm, was added to transcript positions from sequencing-based technologies); and a Yearbook view presenting a paginated array of cell portraits from a single-cell query. Yearbook queries: high *INS* expression (Xenium), randomly selected cells from cluster 5 (Atera), high *IGKC* expression (Visium HD), and high *Col1a2* expression (StrataMap). (B) Celldega visualization of 3D reconstructions generated from Illumina’s StrataMap (pre- commercial) data of a developing whole mouse head across embryonic stages E12.5, E13.5, and E14.5. Left: 3D landscape views showing the spatial organization of cell clusters across developmental time. Right: the same views showing expression of the cartilage marker *Matn1*. (C) Benchmarking Celldega’s performance against spatial visualization tools (Cartoscope, Xenium Web, TissUUmaps, and Vitessce) using Xenium and Atera datasets with transcript totals ranging from 10 million to over 1 billion. Each metric was measured in triplicate; averages are shown. Bars with a jagged top exceeded the 5-minute time limit and are clipped at 300 s; black × markers indicate runs that crashed before completion. (C(i)) Average loading time (solid bar), defined as refreshing the webpage and waiting for the tool to finish loading with no user interaction, and transcript-visualization-enabling time (light bar), defined as the time required to enable transcript visualization for all genes. (C(ii)) Average interactive exploration time, defined as the time required to zoom into a region of interest, wait for rendering to complete, and pan around the region (four times), repeated across three regions, on datasets of increasing size. (C(iii)) Formatted dataset size (GB) for each tool across all four datasets.

### Celldega’s spatial visualization tools provide responsive, flexible, and extensible access to multiple spatial data types

To visualize in physical space, we developed spatial-visualization tools that support common spatial analysis approaches: “Landscape” for multi-scale exploration and “Yearbook” for efficient parallel visualization of queried single-cells and surrounding neighbors (Fig. 2A). Landscape lets users interactively explore large datasets across multiple composable layers, coloring cells by cluster or gene expression, visualizing transcripts at subcellular resolution, overlaying neighborhood polygons, and manually annotating regions of interest (Fig. 2A). As the user pans and zooms, ranked bar charts of cell clusters and genes update on the fly to reflect the local composition of the current neighborhood in the field of view. Yearbook addresses a different task: rather than manually zooming and panning to inspect cells from a population of interest, users supply a list of cells (e.g., representative cells from a cell type) and Yearbook generates a paginated, zoom- synced array of cell "portraits," enabling rapid visual survey of hundreds of cells and their local neighborhoods—useful for confirming cell-type assignments or spotting segmentation artifacts (Fig. 2A, bottom). These tools allow efficient visual exploration of large spatial-omics datasets while easily integrating external analysis results from AnnData.

Celldega’s spatial viewers provide cross-technology and cross-platform visualization of multi-omic spatial data (Fig. 2A). The Landscape view consists of composable image, cell, polygon, transcript, and neighborhood layers. These layers allow Celldega to support both imaging- and sequencing-based spatial transcriptomics data from a wide variety of commercial vendors and non-commercial instruments (Fig. 2A). Celldega similarly supports spatial proteomics visualization, including Xenium with 27-plex protein staining (Supp. Fig. 1A), by extending the number of image layers. The Landscape view also supports 3D visualization using a single-cell-level point-cloud visualization approach (Fig. 2B), which we demonstrate using a 4.5M cell 3D serial section dataset of the developing whole mouse head at three embryonic time points—generated with the pre-commercial Illumina StrataMap platform (Fig. 2A, right; Fig. 2B). This enables highlighting a cluster of cartilage cells and an associated marker gene, *Matn1*, that can be seen increasing in number and intensity across the whole mouse head development (Fig. 2B, right).

To enable a user-friendly, responsive experience, Celldega’s spatial visualizations use a semantic zooming approach, where the visual representations change with scale (Fig. 2A). Biological entities are revealed to the user as they become relevant—streaming data from DegaFiles as needed and utilizing GPU-accelerated rendering. This allows the user to visualize tissues across their natural biological hierarchies—tissue domains and neighborhoods at low magnification, single-cells at intermediate magnification, and cell segmentation polygons and individual transcripts at high magnification (Fig. 2A). This avoids the computational cost and visual clutter of rendering too many objects at once, while guiding users through multi-scale biological structures. We compared Celldega’s features and interactive performance against several existing open-source and commercial tools using multiple large spatial transcriptomics datasets (Fig. 2C; Supp. Tables 1–2; Supp. Fig. 2). Celldega visualized datasets containing millions of cells and over 1 billion transcripts without restricting visualization to preselected genes, sub-sampling transcripts, or aggregating transcripts at high zoom levels (Fig. 2B, Fig. 2C(i-ii)). Its visualization-specific DegaFiles format is substantially smaller than raw instrument data—5–20× smaller than corresponding Xenium outputs and equivalent formatted datasets for open-source tools (Fig. 2C(iii); Supp. Table 2). Across metrics—loading time, transcript visualization enabling time, interactive exploration time, and formatted dataset size—Celldega consistently ranks among the top-performing tools and excels at interactive exploration (Fig. 2. C; Supp. Table 2; Supp. Fig. 2).

### Celldega supports coupled data views

Beyond rendering in physical space, Celldega’s design also supports coupled data and information views—implemented as separate, linked widgets—that connect spatial visualization to downstream analysis outputs (Fig. 1A(i), middle, right; Fig. 1B(i, iii-iv)). First, dimensionality reduction visualizations, such as UMAP, can be a powerful approach to obtain a high-level view of single-cell clustering results and single-cell-level heterogeneity. In addition to visualizing cells in physical space, Landscape can also visualize cells in low-dimensional embedding spaces (Fig. 1B(i)). Landscape includes animated transitions between spatial and UMAP views and lets users interactively explore embedding results by zooming/panning, highlighting cell clusters, and coloring cells by expression level.

Second, spatial-omics workflows produce high-dimensional data (e.g., pseudo-bulked gene expression signatures) whose biological structure is difficult to explore using only flat graph-based clusters or low-dimensional embeddings. Leiden clustering (21) is useful for identifying discrete groups, but it is often necessary to explore nested relationships among coarse and fine-grained cell types, cell states, neighborhoods, and notably gene programs (22–24). To support this, we developed Clustergram (Fig. 1A(i), middle; Fig. 1B(iii)), an interactive visualization for hierarchical biclustering, together with a Matrix class (Fig. 1A(ii-iii), right) for calculating, storing, and linking observation and feature hierarchies, metadata, and clustered matrix values. Building on previous work (23,25) and other interactive heatmap tools (17,26), Clustergram visualizes agglomerative hierarchical bi-clustering results in which both entities (e.g., cells, cell clusters, niches, neighborhoods) and high-dimensional variables (e.g., genes, proteins, populations) are independently clustered and explored through interactive dendrograms. This allows researchers to identify hierarchical structure in their data, overlay prior knowledge as interactive row and column attributes, and link relational views of high-dimensional data to spatial visualizations (Fig. 1B (ii-iii)). The Matrix class supports these workflows by performing preprocessing and clustering while extending AnnData-like organizational principles to include explicit entity types and independent entity/variable hierarchies, improving data provenance and reuse across Celldega views. Alternatively, Clustergram can receive neighborhood-level measurements, linking higher-order spatial structures to high-dimensional data visualization.

Third, exploratory spatial analysis often requires rapid integration of prior biological knowledge (e.g., gene set enrichment analysis), but these workflows are typically performed in separate tools outside the visualization environment. Celldega provides “information” visualizations that integrate biological prior knowledge into visualizations using publicly available APIs (Fig. 1A(i), right; Fig. 1B(iv)). For example, Celldega uses the UniProt API (27) to retrieve gene descriptions for selected genes in the Landscape view, and the Enrichr API (23,28) to perform enrichment analysis on gene sets selected from Clustergram (e.g., clickable dendrogram tree or columns) and visualize these results in the Enrich widget (Fig. 1B(iii-iv)). Together, these tools can guide cell type assignment, cell state identification, and pathway enrichment analysis in the context of spatial and high-dimensional data visualization.

For the rest of the paper, we will show how Celldega can be utilized to facilitate integrative analysis and visualization workflows.

### Neighborhood-level quality control enables segmentation refinement

Accurate cell segmentation is a requirement for properly assigning transcripts to cells and remains a key first step in quality control of spatial samples. Even with improved membrane markers available from the latest generation spatial methods, segmentation remains challenging, particularly in tissues with substantial heterogeneity, where a single segmentation model or algorithm may not perform uniformly across diverse cell types. Segmentation quality is typically evaluated using dataset-wide metrics, such as the proportion of transcripts unassigned to cells, but these global metrics can obscure regional variability in performance across the tissue. Here we demonstrate how Celldega’s notebook-based Landscape visualization and neighborhood methods enable assessment of segmentation quality and integration of improved cell segmentation results in a tissue region with poor performance.

We began by constructing a hextile neighborhood collection and calculating the proportion of transcripts that are unassigned to cells in each hextile across the tissue (Fig. 3A). When visualized in Landscape, this revealed the dermis as a region of the tissue with a particularly high proportion of unassigned transcripts indicating missed or mis-segmented cells (Fig. 3B). Next, we used Celldega’s alpha shape neighborhood method to define a coarse-grained dermis neighborhood using the locations of epidermis- and dermis-associated cell clusters—defining the region to be re-segmented (Fig. 3C). Using cells from the dermis neighborhood, we trained a custom Cellpose2 model (29) to improve segmentation results in this region. This custom model detected more, larger cells and reduced the proportion of unassigned transcripts in the dermis neighborhood—indicative of improved segmentation quality (Fig. 3D). Finally, we used the dermis neighborhood as a mask to merge our Cellpose2 segmentation with the default Xenium segmentation and visualized the updated results using the Landscape visualization (Fig. 3E). By displaying transcripts and cell boundaries together, the zoomed Landscape views (Fig. 3E) provide a qualitative visual confirmation of the improved quality control metrics (Fig. 3D)—notably fewer unassigned transcripts, more numerous cells, and larger cell areas. Thus, Celldega’s Landscape and Neighborhood modules enabled integrated QC, model retraining, and review in a Jupyter notebook environment.

**Fig. 3.**
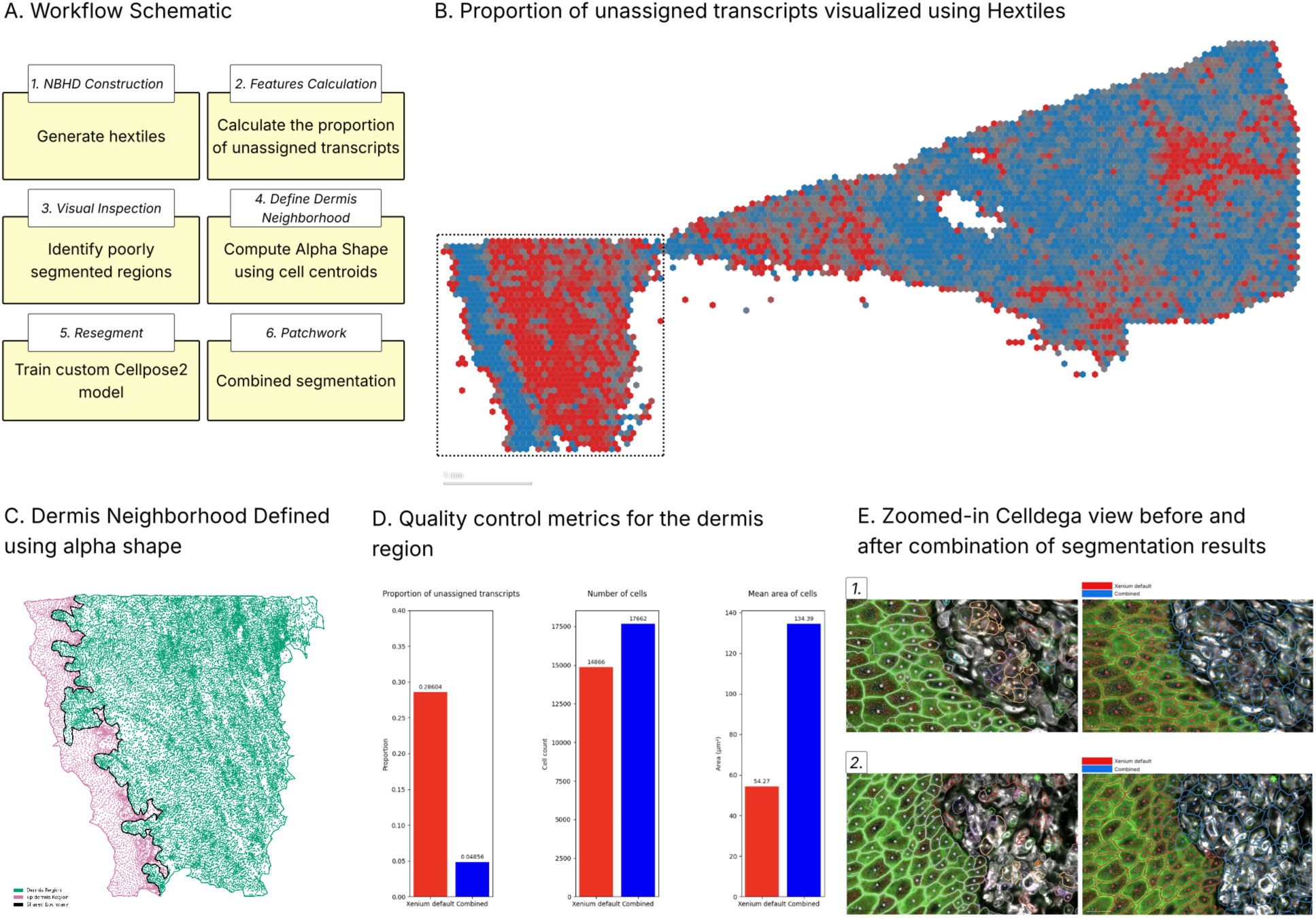
Neighborhood-Level Quality Control Enables Segmentation Refinement. **(A)** Workflow for identifying and correcting a poorly segmented tissue region. **(B)** Celldega Landscape hextile visualization of the proportion of transcripts unassigned to cells enabled visual quality control and revealed a poorly segmented dermal region (shown in red within the dotted outline). **(C)** Dermis (green) and epidermis (pink) alpha shape neighborhoods. **(D)** Bar plots summarizing the dermis neighborhood’s proportion of unassigned transcripts, cell count, and mean cell area. **(E)** Zoomed-in Landscape view before and after integration of Cellpose2 segmentation results.

### Visualization and comparison of spatial domains from external tools using Celldega

Recently a wide variety of methods for defining biological domains in tissues have been developed for spatial transcriptomics data. Benchmarking papers (30,31) provide guidance but still tend to perform apples to apples comparisons of methods solving similar computational problems. However, when interacting with a new dataset it is not always clear what approach to take, and thus a user may seek to compare the results from orthogonal approaches. Here we show how Celldega can be used to perform a meta-analysis of domain identification approaches and identify points of agreement and disagreement (Fig. 4A). To demonstrate this capability, we utilized four algorithms with different approaches: SpaGCN (32), GraphST (33), GASTON (34), and Points2Regions (35) to identify domains in the mouse brain using a publicly available Xenium dataset (Fig. 4B(ii)). We also used Leiden clustering to obtain a definition of tissue domains that are not influenced by spatial information (Fig. 4B(i)). Finally, we turned to Landscape to manually annotate brain regions geometrically—in contrast to the set-based domains produced by most of the algorithms surveyed here (Points2Regions optionally returns domain geometries)—and define a prior-knowledge baseline for mouse brain regions (Fig. 4B(iii)).

**Fig. 4.**
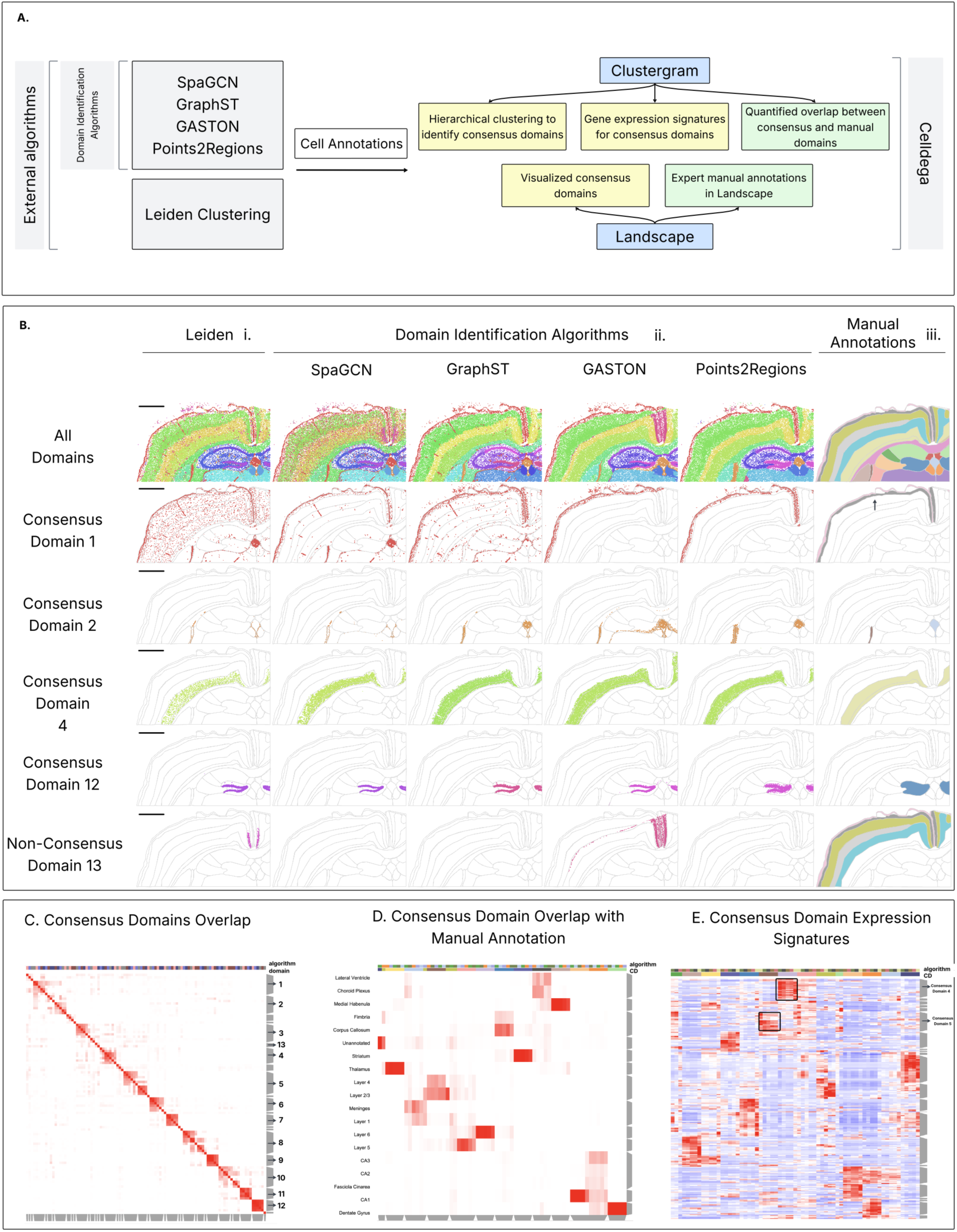
Celldega Enables Cross-Algorithm Domain Identification Comparison and Meta- Analysis. (A) Workflow describing cross-algorithm domain identification comparison using Celldega. (B) Top row: Landscape visualization of Leiden clusters (B(i)), domain identification algorithm domains (B(ii)), and manually annotated regions (B(iii)). Remaining rows: visualization of Consensus Domains and Non-Consensus Domains, overlaid on manual annotation region outlines and aligned with their best-fit manually annotated region(s). Scale bars represent 1 mm. The arrow highlights the manually annotated region that corresponds best to Consensus Domain 1. (C) Clustergram visualization of hierarchical clustering of domain cell-set overlap, identifying cross-algorithm Consensus Domains. (D) Clustergram visualization of hierarchical clustering of domain cell-set overlap (columns) with manual annotation cell sets (rows). (E) Clustergram visualization of Consensus Domain gene expression signatures ordered by Consensus Domain (columns are Consensus Domains, rows are genes). Black outlines indicate gene expression modules associated with Consensus Domains 4 and 5.

We computed pairwise domain–domain similarity using cell-set overlap, clustered the resulting matrix to identify 12 consensus domains and 1 non-consensus domain found across the five algorithms, and visualized them in tissue (Fig. 4B(i-ii); 4C(i); Supp. Fig. 3). Celldega’s linked Clustergram and Landscape views let us directly visualize consensus domains and made points of agreement and disagreement immediately apparent (Fig. 4B(i- ii), 4C(i)): all algorithms recovered major structures (cortical layers 5/6, dentate gyrus, CA1, CA3), while finer distinctions—such as separating layer 1 from the meninges—were captured inconsistently (consensus domain 1, Fig. 4B(iii); Supp. Fig. 3). The framework also surfaced method-specific behavior, including GraphST’s detection of the transcriptionally distinct proliferative sublayer in the dentate gyrus (Fig. 4B(ii), Consensus Domain 12), GASTON’s strong enforcement of spatial continuity (Fig. 4B(ii), Non-Consensus Domain 13), and Leiden’s mixed-layer domains (Supp. Fig. 3(i)).

With this initial analysis carried out, we wanted to identify manually annotated regions that consistently agreed/disagreed with algorithmically defined domains. We generated a Clustergram to visualize the cell-set overlap between the cells that are contained within manually annotated domains and the algorithmically derived consensus domains (Fig. 4D).

Consistent with the comparisons described above, the Clustergram showed strong and specific overlap between consensus domains and major manually defined anatomical regions, including the striatum, thalamus, dentate gyrus, CA1, and deep cortical layers. In contrast, upper cortical layers (layer 1, 2/3, and 4) showed less agreement with algorithmically defined domains and were often distributed across more than one consensus domain with neighboring layers showing partial overlap. These differences were primarily confined to adjacent cortical regions rather than unrelated structures, indicating that the algorithms tend to generally preserve overall anatomical organization while differing in how finely they partition laminar boundaries (Fig. 4D).

Finally, we calculated consensus domain gene expression signatures to help interpret biological differences between domain definitions. We explored these data using a linked Clustergram and Enrich widget for on-the-fly enrichment analysis of gene clusters and programs (Fig. 4E). We identified that the consensus domain 4 exhibits a robust and cross-algorithm enrichment of layer 5/6 neuronal markers, including Fezf2 (36) and Tle4 (37), supported by CellMarker_2024 enrichment analysis. The same gene set showed relatively low expression in neighboring consensus domains (blue regions), indicating that the deep-layer transcriptional signature is spatially restricted and sharply delineated, rather than part of a gradual gradient. In contrast, consensus domain 5 (Supp. Fig. 3) displayed a transcriptional profile characteristic of superficial cortical layers, with enrichment of genes such as Cux2 and Rorb, which are commonly associated with Layer 2/3 and Layer 4 neuronal populations (Supp. Fig. 3) (38–40). Permanent links to enrichment results are provided in the Data Availability section. Taken together, Celldega enabled a meta-analysis of domain identification algorithms through a flexible multimodal visualization framework.

### Celldega integrates spatial transcriptomic and proteomic data to resolve T cell functional states in human renal cell carcinoma

Spatial datasets are becoming increasingly multimodal, with most technologies at the very least supporting paired HCE imaging, and an increasing number enabling paired immunofluorescence staining. To test Celldega’s suitability for multi-omic analysis, we resolved T cell subtypes and states in a publicly available human renal cell carcinoma Xenium dataset with 405 genes and 27 immune-focused proteins (Fig. 5). We first identified 12 coarse-grained cell clusters in gene expression space and utilized linked Landscape, Clustergram, and Enrich widgets to manually annotate cell type using a combination of marker gene expression, enrichment analysis, and interactive exploration of cells in UMAP and spatial views (Fig. 5A) (see Methods). These clusters grouped into four larger meta- clusters (MCs; Fig. 5A(iii)), corresponding to the myeloid compartment (MC-1), stroma/vasculature compartment (MC-2), lymphoid compartment (MC-3), and tumor/neoplastic compartment (MC-4).

**Fig. 5.**
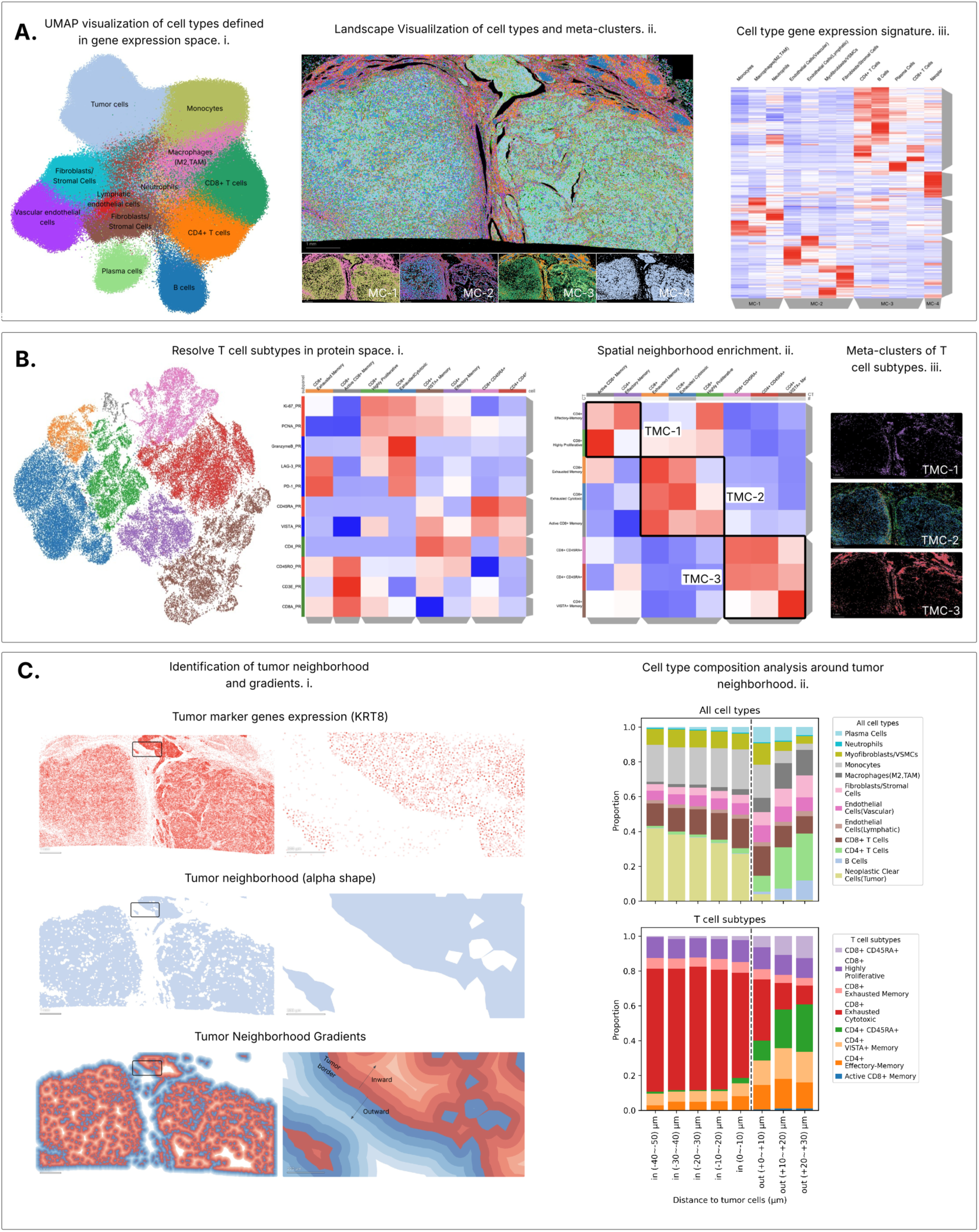
Integrated Single-Cell Spatial Transcriptomic and Proteomic Analysis Resolves T Cell Functional States in Human Renal Cell Carcinoma. **(A)** Single-cell transcriptional profiling and cell type identification. **(A(i))** UMAP visualization of the gene expression space embedding, colored by cell type obtained through manual annotation of Leiden clusters. **(A(ii))** Landscape visualization of cell types (top) and meta-clusters (MC) (bottom): MC-1, infiltrating myeloid compartment; MC-2, stroma/vasculature compartment; MC-3, lymphoid compartment; MC-4, tumor/neoplastic compartment. Scale bar represents 1 mm. **(A(iii))** Clustergram visualization of cell type gene expression signatures. Meta-clusters are highlighted in the column dendrogram. **(B)** T cell proteomic profiling and functional state identification. **(B(i))** UMAP visualization of the protein expression space embedding, colored by T cell subtype and functional state obtained through manual annotation of Leiden clusters. Clustergram shows proteins relevant to T cell function (rows) and functional T cell states (columns). **(B(ii))** Hierarchical clustering of neighborhood enrichment analysis results, revealing three T cell meta-clusters (TMC): TMC-1 (active effector), TMC-2 (CD8+ exhaustion), and TMC-3 (resting/naive-like). **(B(iii))** Landscape visualization of T cell neighborhood-enrichment-based meta-clusters (TMC). **(C)** Gradient cell type composition analysis around the tumor neighborhood. **(C(i))** Top row, tumor marker gene expression; middle row, tumor neighborhood (alpha shape); bottom row, gradient neighborhoods. **(C(ii))** Cell type composition (top) and T cell functional state composition (bottom) as a function of distance from the tumor neighborhood.

T cells were selected from the lymphoid compartment and refined using canonical protein markers to enrich for CD3E-positive T cells while excluding B cells, myeloid cells, and epithelial/tumor cells (+CD3E, −CD20, −CD68, −PanCK; see Methods). We then utilized linked UMAP and Clustergram views to visualize cells clustered in protein space (an 11- dimensional T-cell-relevant protein space) and manually annotate eight T cell subtypes and functional states (Fig. 5B(i)). Next, we used Clustergram to explore how T cell subtypes co- localized in the tissue. We performed neighborhood-enrichment analysis using Squidpy (7) and visualized the results as a Clustergram (Fig. 5B(ii)). Hierarchical clustering of these spatial-enrichment profiles identified three spatial T cell neighborhoods (TNs): TN-1, active effector neighborhood (CD4+ effector-memory with active CD8+ memory); TN-2, exhausted/proliferative CD8+ neighborhood (CD8+ exhausted memory, CD8+ exhausted cytotoxic, and CD8+ highly proliferative); and TN-3, resting/naive-like neighborhood (CD8+ CD45RA+, CD4+ CD45RA+, and CD4+ VISTA+ memory) (Fig. 5B(ii-iii)).

To further characterize the spatial architecture of the tumor microenvironment (TME), we analyzed cellular composition as a function of distance from the tumor boundary. Tumor cells were identified using carcinoma and clear-cell-associated marker genes, including KRT8, MKI67, MET, and CDH2 (Fig. 5C(i)). We then defined the tumor neighborhood using Celldega’s alpha-shape method and generated concentric 10 μm regions extending inward and outward from the tumor boundary (see Methods). This analysis revealed a clear transition in cellular composition across the tumor boundary (Fig. 5C(ii)). Regions outside the tumor neighborhood were enriched for stromal and immune populations, including fibroblasts, B cells, CD4+ T cells, and CD4+ VISTA+ memory T cells. Moving inward across the tumor boundary, the proportion of tumor cells increased, accompanied by a shift in T cell subtype composition toward CD8+ exhausted cytotoxic cells. These results show how Celldega can integrate protein-defined immune states with spatial region analysis to characterize immune organization around tumor boundaries.

## Discussion

Celldega demonstrates a scalable framework for integrating high-performance visualization with spatial transcriptomics analysis. In the examples presented here, Celldega is used across a wide variety of spatial data types and throughout the spatial-analysis lifecycle— quality control, exploratory analysis, annotation, algorithm comparison, biological interpretation, and dissemination. Through these examples we have highlighted a recurring theme: that many spatial questions require representations and analyses of higher-order biological entities that reflect biological hierarchies in tissue organization. Celldega supports this with lightweight data structures and analysis methods in which neighborhoods are treated as first-class objects coupled directly to the visualization layer, alongside biclustering methods that enable exploration of cell and gene hierarchies. These methods are meant to complement rather than replace established single-cell and spatial toolkits such as Scanpy (41) and Squidpy . While several tools exist in the field which seek to address this gap, Celldega’s key differentiators are its flexibility and responsiveness.

With regards to responsiveness, user interface research (42,43) has shown that slow response times measurably degrade usability, disrupting users’ flow and attention during interactive tasks. Celldega’s Landscape visualization maintains response times in the one second range, even for datasets with >1B transcripts, allowing researchers to explore datasets fluidly rather than waiting on the interface (example video: github.com/broadinstitute/celldega#celldega-demo-video). This responsiveness is designed to hold as datasets and cohorts grow: Celldega’s tiling approach scales naturally with data size, and its multi-scale semantic zooming keeps rendering performance stable as spatial transcriptomics datasets continue to increase in scale. Flexibility comes across in three ways.

First, Celldega is designed primarily as a visualization tool to be used in combination with external spatial analysis workflows: its notebook-first design and modular, high-performance visualizations let researchers integrate Celldega directly into spatial analysis notebooks, whereas other tools often require stepping outside the notebook or are meant to be used only after analyses are complete. Thus, Celldega can be closely integrated with virtually any spatial analysis without requiring further redevelopment.

Second, the minimal additional data types and structures allow simple extension to new techniques. Compact visualization specific DegaFiles are the key Celldega specific format. We currently support the most common spatial transcriptomics technologies and intend to support more in future development. However, the already developed readers for SpatialData mean DegaFiles can inherit the ease of import built out by the Spatial Data consortium. To emphasize this input format flexibility, we have shown that Celldega even extends easily to non-spatial data such as single cell sequencing and perturb-seq (Supp Fig 4A).

Third, building Celldega on the Jupyter Widgets framework lets it take full advantage of the notebook ecosystem—and, more recently, the Marimo/Molab Python notebook ecosystem. This enables researchers to embed interactive, linked visualizations into executable notebooks and public galleries. At broadinstitute.github.io/celldega/examples, users can browse these galleries, open executable notebooks on Google Colab and Molab, and find tutorial videos covering Celldega’s features. We built Celldega’s visualizations as Jupyter Widgets specifically for their modularity and extensibility. Although the Widgets framework has existed for over a decade, building new widgets has traditionally been difficult, slowing adoption. However, recent developments—notably anywidget and AI-assisted software development—have substantially lowered this barrier. To support third-party development, we provide API-compatibility guidelines in our documentation. Following these ourselves, we built a custom geospatial widget—linked to a Clustergram—for exploring bike-share stations clustered by destination similarity and visualizing their associated geospatial neighborhoods, and released it as a standalone pip-installable package (Supp. Fig. 4B; Supplementary Note).

The design choices above have downstream implications. Celldega runs well in cloud-hosted environments such as Terra.bio, Manifold.ai, Code Ocean, and Saturn Cloud. Because DegaFiles are compact, they can be hosted on free repositories such as GitHub and Hugging Face or on public cloud storage and served directly into notebooks or web pages, supporting open science and FAIR data principles (44). All interactive examples in Celldega’s documentation are powered this way. Celldega is also compatible with free web-based notebook services (Google Colab and Molab), can be organized into shareable public galleries, and can be embedded into any webpage or web app by importing it as an ECMAScript module—letting interactive visualizations live directly in project websites, data portals, or custom applications.

Currently Celldega has several limitations that we will address with future developments: scaling workflows toward large cohort-scale spatial studies, developing additional workflows for multi-modal datasets, and developing additional "information" visualizations (e.g. AI assisted interpretation). Alongside this, we will continue to expand tutorials, documentation, and example galleries, and improve interoperability with open-source tools, to facilitate adoption by both computational and bench scientists. Taken together, Celldega establishes an open, scalable, and extensible framework in which visualization is not only a final presentation step but an active component of spatial-omics analysis and biological discovery.

## Methods

### Samples and datasets

#### Public Datasets

Public spatial transcriptomics datasets used in this study were obtained from the 10x Genomics and Bruker Spatial Biology websites. These included Xenium In Situ Gene Expression, Xenium In Situ Gene and Protein Expression, and Visium HD datasets used for the examples shown in Figures: 1B, 2A, 3, 4, and 5; Supp. Figs. 1A, 2, and 3. We used a public Human colon cancer (whole transcriptome) CosMx dataset (Supp. Fig. 1B). The CRISPR activation perturbation dataset was obtained from the Pertpy library; this dataset was originally described by Norman et al. (45) and consists of 111,255 K562 single-cell transcriptomes across 287 single-gene and gene-pair perturbations (Supp. Fig. 4A).

#### Illumina StrataMap Pre-commercial dataset

We obtained developing whole mouse head data using Illumina’s StrataMap (pre-commercial assay) in collaboration with Paola Arlotta’s laboratory and the Illumina Inc. Research and Development team. Whole mouse heads of C57BL/6 mouse embryos at key stages of cortex formation (E12.5, E13.5, and E14.5) were frozen in OCT using an isopentane liquid nitrogen bath. Specimens from each stage were serially sectioned along the antero- posterior axis, from forebrain through the head. A 10-micron section was collected every 200 microns (10 sections total) and placed on the assay substrate. Sections were fixed in methanol, stained with hematoxylin and eosin (HCE), and imaged in brightfield using a Keyence BZ-X810 microscope. Slides were then processed following a developmental pre-release version of the StrataMap assay workflow. This resulted in over 4 million single cells, which were clustered across all time points (using a Scanpy single-cell clustering workflow). Celldega was used to visualize the spatial distribution of cell types in 3D after manual alignment of 2D slices (Fig 2B).

### Visualizations

Celldega’s visualizations are built in JavaScript and use deck.gl for WebGL-based rendering, D3.js for interactive control panels, and a Parquet/Arrow approach for efficient browser-side rendering of spatial data. Celldega uses a custom observable store approach for state- management to coordinate visualization state across layers and linked widgets.

#### Landscape

Landscape renders spatial-omics data as composable deck.gl layers, including images, cell centroids, cell segmentation geometries, transcript coordinates, and neighborhood annotations. Spatial features are loaded from DegaFiles using viewport-dependent tile loading, allowing the browser to retrieve only the data required for the current view. Interactive controls manage layer visibility, cell-annotation filters, transcript-gene filters, and viewport-level summaries. Cell and transcript counts are recomputed for the visible viewport and displayed as live bar plots in the control panel. Cell annotations and numerical metadata can be provided through an AnnData object and transferred from adata.obs to the front end using widget traitlets; embedding coordinates, such as UMAP coordinates, can also be provided through AnnData and linked to the same cell identifiers. Neighborhood collection geometries are provided as GeoPandas GeoDataFrames or NeighborhoodCollections and synchronized to the front end using widget traitlets. Manual spatial annotations are implemented using editable deck.gl layers and synchronized back to Python as GeoPandas-compatible geometries for downstream analysis.

#### Yearbook

Yearbook is implemented as a multi-view spatial visualization for inspecting queried cells and their local neighborhoods in parallel. Yearbook reuses the same layer and control-panel architecture as Landscape but renders many synchronized cell-centered viewports rather than a single contiguous spatial viewport. The front end uses deck.gl with shared rendering layers, multiple synchronized viewports, and synchronized zoom state to display a grid of cell portraits. To reduce data transfer, Yearbook performs discontiguous tile loading, retrieving only the image, cell, boundary, and transcript tiles required for the currently displayed set of queried cells. In addition to the standard Landscape layer controls, Yearbook adds a lightweight front end query interface for selecting cells by identifier, categorical annotation, or gene expression driven criteria, together with pagination controls for reviewing queried cells across multiple pages.

#### Clustergram

Clustergram widgets are initialized from Celldega Matrix objects. Matrix values, metadata, and linkage information are transferred from Python to the JavaScript front end using Parquet-encoded widget traitlets. The Clustergram visualization front end renders high- dimensional data as an interactive heatmap with row and column attributes and interactive dendrograms—rendered as dendrogram clusters formed by cutting the dendrogram tree at an interactively defined height. The visualization is built using deck.gl and D3.js and is implemented using multiple synchronized orthographic viewports, allowing the matrix, attributes, and dendrograms to maintain coordinated zoom and pan states and to support interactive animated reordering. Custom zoom and pan constraints restrict navigation to valid matrix regions and support progressive exploration of large matrices. Matrix values, metadata, clustering results, selections (e.g., dendrogram clicks), interaction events, and user annotations are accessible through the widget API, enabling reuse across linked Celldega visualizations.

#### Enrichr and UniProt integrations

Celldega integrates external biological knowledge through lightweight information- visualization widgets (e.g., Enrich). For gene set enrichment, selected gene lists are submitted to the Enrichr API, and returned terms, significance values, and scores are parsed and displayed as an interactive bar visualization. Gene annotation information is retrieved from the UniProt API and displayed with selected genes to support interpretation during exploratory analysis. Use of these external services is subject to their respective terms of service and licenses.

#### Notebook integration

Celldega’s Python visualizations are implemented as Jupyter widgets using AnyWidget (46), enabling bidirectional communication between Python analysis objects and JavaScript visualization components. Widgets are initialized with data passed through arguments (e.g., the DegaFiles URL). After initialization spatial visualizations load tiled data directly on the front end by making RESTful requests to hosted DegaFiles, minimizing the amount of data transferred through the notebook kernel. DegaFiles can be hosted locally using a small local server (provided by Celldega) or on public hosting services (e.g. GitHub). Optional AnnData objects can be provided to the Python API, allowing cell annotations and numerical metadata from adata.obs to be transferred to the front end for visualization. Bidirectional widget communication is also used to return user-generated annotations from the front end to Python, including spatial annotations from Landscape and categorical annotations from Clustergram Celldega widgets are compatible with: Jupyter Notebook, Jupyter Lab, Google Colab, Marimo, and Molab. A preconfigured Celldega environment is available to Broad Institute users through Manifold.ai.

### DegaFiles

DegaFiles are visualization-specific, self-contained directories generated from raw spatial- omics outputs using Celldega’s preprocessing module. Each set of DegaFiles contains the data required for interactive rendering, including multiscale image tiles, spatial feature tiles, cell-boundary geometries, transcript coordinates, cell-by-gene expression data, and metadata. Images are stored as multiscale WebP pyramids, while spatial features and expression matrices are stored in Apache Parquet format. Spatial features are tiled to support viewport-dependent loading, and expression matrices can be stored either as one Parquet file per gene or as chunked Parquet files using Parquet row groups to reduce file counts while preserving selective access. A manifest records dataset configuration, image channels, tile size, file locations, and chunking strategy. Additional details are provided in the online Celldega documentation.

#### Matrix Class

High-dimensional data that will be hierarchically clustered and visualized in the Clustergram is stored in the Matrix class, which stores matrix values, row and column metadata, and hierarchical biclustering results. The Matrix class accepts tabular data or AnnData objects and supports preprocessing steps such as normalization, filtering, and aggregation. Hierarchical biclustering is performed using SciPy, with support for variable distance metrics and linkage methods. The resulting row and column linkage matrices are stored with the matrix values and metadata. These data are transferred to Clustergram widgets using Parquet-encoded widget traitlets for efficient exchange between Python and JavaScript. Matrix values, metadata, and clustering results are accessible through the Python API and can be reused across Celldega visualizations.

#### Neighborhood Class

Celldega represents neighborhoods using the NeighborhoodCollection (nbhd) class. A neighborhood is represented as a polygon or multipolygon geometry, and neighborhood collections are stored as MuData objects with GeoDataFrame geometries. Neighborhood collections can be generated from AnnData coordinates, imported from external spatial domain identification algorithms, or created from manual Landscape annotations.

The NeighborhoodCollection class implements three geometry-generation methods. Hextile neighborhoods are generated by tiling the spatial extent of an AnnData object with non-overlapping hexagons using a user-defined diameter in microns. Alpha shape neighborhoods are constructed from sets of cell coordinates using the alpha_shape function in the libpysal.cg module. Celldega calculates alpha shapes for cells from each categorical value for a given attribute (e.g., all Leiden cluster categories). Gradient neighborhoods are generated from a given neighborhood by geometric buffering—producing concentric inward and/or outward rings at a user-defined step size—and is calculated using GeoPandas.

The NBHD class computes neighborhood-level feature spaces by intersecting neighborhood geometries with cells, transcripts, or image-derived measurements. These include neighborhood-by-gene matrices from cell-level expression or raw transcript counts, neighborhood-by-population matrices from categorical cell labels, neighborhood-by-image summaries, and neighborhood-level metadata such as cell counts, transcript counts, proportion of transcripts not assigned to cells, area, and perimeter. Pairwise neighborhood relationships can also be computed using overlap metrics such as intersection-over-union and border metrics such as shared border length. Resulting matrices can be passed to the Celldega Matrix class for clustering and visualization.

#### Visualization Benchmarking

Celldega (v0.16.0) was benchmarked against five browser-based spatial transcriptomics visualization tools: Cartoscope (18), Xenium Web, TissUUmaps (v3.2.1.14) (13), Vitessce (v3.8.4) (17), and the Allen Institute Spatial Explorer (19). Given differences in available datasets, the benchmark is reported in two parts: a primary comparison among Celldega, Cartoscope, Xenium Web, TissUUmaps, and Vitessce (Fig. 2C; Supp. Fig. 2B), and a separate comparison between Celldega and the Allen Institute Spatial Explorer (Supp. Fig. 2C). All benchmarks were run on a 2023 14-inch MacBook Pro with an Apple M2 Max chip (12-core: 8 performance + 4 efficiency), 32 GB RAM, running macOS 26.6 (Tahoe). All tools, including the Jupyter-based tools accessed via localhost, were run in Google Chrome (version 151.0.7922.77) over the Broad Institute’s internal Wi-Fi network. Interactive exploration was performed using an attached external display.

For the primary benchmark, four datasets of increasing scale were used: (1) ∼150k cells / 10M transcripts (Xenium), (2) ∼400k cells / 120M transcripts (Xenium), (3) ∼700k cells / 200M transcripts (Xenium), and (4) ∼720k cells / 1 billion transcripts (Atera). Each dataset includes four imaging channels. Where possible, all visualization layers supported by Celldega were replicated across tools: cell centroids, cell boundary geometries, transcripts, and images.

For each tool, four metrics were recorded per dataset, with each measurement taken three times and averaged. The first metric was formatted data size, the on-disk storage footprint of each tool’s prepared data (Fig. 2C(iii)). Each tool required its own input format. Celldega uses DegaFiles, with storage footprint measured by checking the directory size. The original instrument output data sizes for Xenium and Atera datasets were obtained by downloading the data and checking the directory size, to serve as a baseline. For Cartoscope, dataset sizes were obtained by querying the public Cartostore S3 bucket via the AWS CLI with anonymous access (--no-sign-request). Dataset directories were identified from the appropriate batch and collection prefixes, and total directory sizes were obtained using recursive S3 listings with summary statistics (aws s3 ls --recursive --summarize). Reported sizes represented the full storage occupied by each Cartostore dataset directory, including visualization-specific assets such as PMTiles, image pyramids, metadata, and configuration files, rather than the original instrument outputs. For Xenium Web, the size of the smaller Xenium Output Bundle listed on the official 10x Genomics website was used; the desktop Xenium Explorer application was not included in this benchmark. For TissUUmaps, transcript- and cell-level AnnData (.h5ad) files were generated rather than CSV files, following the tool’s H5AD-based loading pathway for faster marker display. For Vitessce, a SpatialData object in Zarr format was created following a dedicated example notebook from the project’s GitHub repository.

The second metric was average loading time, defined as the time from initiating the tool to completion of initial rendering (Fig. 2C(i); Supp. Fig. 2B(i)). Celldega, TissUUmaps, and Vitessce are Jupyter-based; loading time was measured from running the widget cell to the tool finishing rendering. For Cartoscope and Xenium Web, both web-based, loading time was measured from page load to reaching a fully rendered initial state. No user interaction time was included during this phase. The third metric was average transcript visualization enabling time, defined as the time required for a user to enable the visualization of transcripts from all genes—that is, to enable the display of all transcripts when zooming into a region—plus the time for the tool to finish rendering transcripts (Fig. 2C(i); Supp. Fig. 2B(ii)). Celldega, TissUUmaps, and Vitessce display all transcripts in a region automatically when zoomed in, requiring no additional user action. For Cartoscope, a CSV file containing all gene names must be uploaded to enable this view. For Xenium Web, an option to display all genes was selected from a dropdown; for the three largest datasets, Xenium Web did not display all transcripts even after doing so, suggesting the data may be subsampled.

The fourth metric was average interactive exploration time, defined as the time for a user to zoom into a region, wait for all visible layers to fully render (cell boundary geometries, cell centroids, transcripts, and images), then pan in all four directions (Fig. 2C (ii); Supp. Fig. 2B(iii)). This cycle was repeated across three distinct regions per dataset. In cases where transcript visualization enabling time already reached the 5-minute time limit for every run for a specific tool, the full three-region interactive exploration cycle was not repeated three times but performed once for record-keeping of the behavior, and the exploration time for such runs was hence clipped at the 300-second limit, consistent with the treatment described above.

The Allen Institute Spatial Explorer was benchmarked separately using the two Atera datasets available through the tool: one with approximately 40k cells and 161M transcripts, and one with approximately 300k cells and 1,056M transcripts. These transcript counts are estimated from per-cell means reported by 10x Genomics segmentation (161M ≈ 42,851 cells × 3,757 transcripts/cell; 1,056M ≈ 320,885 cells × 3,291 transcripts/cell). Two Celldega datasets of roughly comparable scale were selected for this comparison: ∼700k cells / 200M transcripts (Xenium) and ∼720k cells / 1 billion transcripts (Atera). The same four metrics were evaluated, with two tool-specific considerations. First, the Allen Institute Spatial Explorer does not make its underlying data publicly available, so a formatted dataset size could not be obtained and is excluded from the formatted dataset size comparison. Second, for transcript visualization enabling time, the Allen Institute Spatial Explorer requires users to manually enter gene names. We limited ourselves to visualizing 50 genes based on the manual search required to activate a gene, resulting in a substantially reduced transcript count of approximately 5M and 20M transcripts for the smaller and larger datasets respectively, compared to the full 161M and 1,056M transcripts present in each (Supp. Fig. 2C).

A 5-minute (300-second) time limit was imposed per phase. In figures, bars that reached this limit are clipped at 300 seconds and indicated with diagonal slash marks (///) at the bar top (Fig. 2C(i-ii); Supp. Fig. 2B-C). Where loading and transcript visualization enabling time are presented as a combined stacked bar, the clip is applied to the combined total (Fig. 2C(i)).

Runs in which the tool crashed before completing a phase are marked with a black × marker. Two crash scenarios were distinguished: (1.) the tool crashed during a prior phase and never reached the phase being measured — for example, TissUUmaps crashed during loading at the 120 M and 200 M Xenium datasets, meaning transcript visualization enabling and exploration times could not be recorded for those conditions (Supp. Table 2); and (2.) the tool crashed during the phase itself after a time had already been recorded — as observed for Xenium Web interactive exploration at the 200 M Xenium dataset, where all three runs produced times but the tool crashed before the session concluded (Supp. Table 2). In the latter case, the recorded time is retained in the average and a bar is displayed, with an × marker above it to flag the crash instability. Notably, TissUUmaps at the 1 B Atera dataset appeared to load but seemed functionally stalled: zooming in revealed only cell centroids, with no cell boundary geometries or transcript coordinates rendered, and the marker loading bar never reached completion.

In cases where runs within a phase showed mixed outcomes — some exceeding the time limit and others crashing — only runs that yielded a valid recorded time contributed to the reported average. This occurred for Xenium Web transcript visualization enabling at the 1 B dataset, where two runs crashed during this phase before producing a completed record time, and only the one run that reached the 5-minute limit contributed to the reported average (Supp. Table 2).

For the Allen Institute Spatial Explorer comparison, an asterisk (*) is shown above bars in the exploration time figure to highlight that this tool was evaluated on a subsampled gene panel of 50 genes, as described above, rather than the full transcript complement used for all other tools and conditions (Supp. Fig. 2C(iii)).

#### Neighborhood-level quality control enables segmentation refinement

Cell segmentation quality in a public Xenium skin melanoma dataset was assessed using the Celldega Neighborhood class. Spatial neighborhoods were generated using hexagonal tiling. For each hextile, segmentation quality was summarized as the proportion of transcripts that are unassigned to cells, calculated from the original transcript assignment provided by the instrument. Hextiles with high unassigned-transcript proportions corresponded to the dermis, indicating reduced segmentation performance in this region (shown in blue within the black-outlined box; Fig. 3B).

To improve segmentation in this region, dermal cells were re-segmented using a custom-trained Cellpose2 model, while epidermal cells retained the original Xenium segmentation. Cellpose2 custom model training followed an iterative curation approach in the Cellpose graphical user interface (GUI): preliminary segmentation masks were first generated using the pretrained Cellpose "cyto3" model applied to the morphology focus image, which contains four channels (DAPI, boundary, interior-RNA, and interior-protein). These preliminary annotations were manually reviewed and corrected within the GUI across four non-overlapping subsections of the tissue image, constituting the full training set. The custom model was then trained using the boundary channel as the primary channel and the DAPI channel as the secondary channel, with "cyto3" as the pretrained base. During inference, a flow threshold of 0.8 and a cell probability threshold of −1 were applied.

The dermis–epidermis boundary was defined by computing an alpha shape from the cell centroids of both epidermis-specific and dermis-specific cell clusters, generating a spatial mask used to separate the two regions (Fig. 3C). The final patchwork segmentation combined epidermal cells from the original Xenium segmentation with dermal cells from the custom Cellpose2 segmentation (Fig. 3D).

Combining segmentation outputs introduced overlap conflicts near the dermis–epidermis boundary. To resolve these conflicts, we computed the intersection over minimum area (IoMA) for all overlapping cell pairs across the two segmentation sources. For overlapping pairs with IoMA > 0.2, the larger cell by area was retained and the smaller overlapping cell was removed. Cells with IoMA ≤ 0.2 were retained without modification.

#### Visualization and comparison of spatial domains from external tools using Celldega

A publicly available mouse brain Xenium dataset was used to compare spatial-domain assignments generated by external algorithms. Spatial domains were computed using SpaGCN (32), GraphST (33), GASTON (34), and Points2Regions (35). Each method produced one domain label per cell. Leiden clustering was also performed on the gene expression matrix using Scanpy to generate a non-spatial clustering baseline. Parameters for each method were selected to produce a comparable number of domains or clusters (n = 20).

Domain labels from each method were imported into Celldega and visualized in Landscape together with spatial gene expression patterns and fluorescence images. Manual anatomical regions were annotated using the Landscape interface while inspecting spatial gene expression and image data (Fig. 4B(iii)). These annotations were stored as categorical cell labels and visualized alongside algorithm-derived domain assignments (Fig. 4B).

Pairwise similarity between domains was calculated using cell-set overlap. For each pair of domains, similarity was defined as the intersection-over-union (IoU) between the sets of cells assigned to each domain. The resulting domain-by-domain similarity matrix was hierarchically clustered to identify groups of highly overlapping domains across methods (Fig. 4C). These groups were summarized as consensus domains, defined as clusters of algorithm-derived domains that shared a substantial fraction of assigned cells.

Overlap between manually annotated anatomical regions and consensus domains was visualized using Clustergram (Fig. 4D). Rows corresponded to manual anatomical regions, columns corresponded to consensus domains, and matrix values represented cell-set overlap. Gene-expression signatures were computed for each consensus domain and visualized as a hierarchically clustered domain-by-gene matrix using Clustergram (Fig. 4E). Selected gene sets corresponding to Consensus Domains 4 and 5 were submitted to the Enrichr API through the Celldega Enrich widget to retrieve enriched biological annotations.

#### Spatial multi-omic analysis of human renal cell carcinoma using Celldega

A public 10x Genomics Xenium renal cell carcinoma dataset with 405 gene targets and 27 protein markers was used for spatial multi-omic analysis. Initial quality control retained cells with at least 10 counts and genes detected in at least five cells. High-level cell types were identified by performing Leiden clustering on the gene-expression matrix using Scanpy, resulting in 12 clusters. Clusters were manually annotated into major cell types using known marker genes and the Celldega Enrich widget.

T cells were selected from the annotated CD4+ and CD8+ T cell populations and further refined using protein-marker expression. High-confidence T cells were identified by manual gating for CD3E-positive cells while excluding cells positive for mutually exclusive lineage markers: CD20 for B cells, CD68 for myeloid cells, and PanCK for epithelial or tumor cells. Log-transformed marker thresholds were set to 4.8 for CD3E, 1.0 for CD20, 2.5 for CD68, and 1.0 for PanCK (Fig. 5B). The resulting T cell population was clustered in 11-dimensional protein space using Leiden clustering, producing eight T cell subtypes or functional states.

Spatial relationships among T cell subtypes were quantified using Squidpy neighborhood-enrichment analysis. Spatial neighbors were computed using sq.gr.spatial_neighbors, and neighborhood enrichment was calculated using sq.gr.nhood_enrichment. The resulting enrichment matrix was visualized using Clustergram (Fig. 5B).

To analyze cellular composition as a function of distance from the tumor boundary, tumor cells were used to define a tumor neighborhood using Celldega’s alpha shape method. Concentric spatial rings were then generated from this tumor region with a bin width of 10 μm and a maximum distance of 50 μm. Cell-type composition summaries were calculated for each spatial ring (Fig. 5C).

#### Public galleries and embedded visualizations

Interactive Celldega examples for the documentation and public galleries were generated from notebooks and exported as standalone web artifacts. For each example dataset, DegaFiles were generated and hosted on a public repository or cloud-accessible storage location. Example notebooks then instantiated the relevant Celldega widgets, including Landscape, Clustergram, Yearbook, and Enrich, using URLs of public DegaFiles. These notebooks were used both as executable analysis examples and as sources for rendered documentation pages.

Standalone gallery views were generated by embedding Celldega’s JavaScript visualizations into static webpages. Because DegaFiles are loaded directly by the browser, these gallery pages do not require a dedicated backend server after preprocessing. The resulting pages can be hosted through static-site infrastructure, including GitHub Pages, documentation websites, or public cloud storage. For examples intended for interactive notebooks, we also prepared Google Colab-compatible notebooks that load the same public DegaFiles and reproduce the corresponding Celldega visualizations.

## Data availability

Except for the developing whole mouse head data, which was acquired via a pre-release StrataMap assay and will be made public upon peer-reviewed publication, all other datasets used in this study are publicly available and summarized in this GitHub repository (https://github.com/broadinstitute/Celldega_Public_Dataset_Directory). Original instrument and formatted visualization tool-specific datasets used in the performance benchmark along with screen recordings can be found here: https://huggingface.co/datasets/broadinstitute/Celldega_Visualization_Benchmarking. Enrichr results for Consensus Domain 4 gene cluster are available here: https://maayanlab.cloud/Enrichr/enrich?dataset=0f7270960f88c89ec8f19b4af23666a5. Enrichr results for Consensus Domain 5 gene cluster are available here: https://maayanlab.cloud/Enrichr/enrich?dataset=614e40b5b3a0c69e529f2e91f996e255.

## Code availability

The Celldega source code is available via GitHub (https://github.com/broadinstitute/celldega) under the Broad Institute Academic Software License (https://github.com/broadinstitute/celldega/blob/main/LICENSE.txt). Further documentation, tutorials, gallery, and examples notebooks are available at https://broadinstitute.github.io/celldega/.

## Supporting information

Supplementary_Table_1_Features

Supplementary_Table_2_Benchmarking

## Acknowledgements

We thank Paola Arlotta and Daniela Di Bella for contributing whole mouse head samples and collaborating on data analysis. We thank Illumina Inc. for their collaboration and for providing access to the pre-commercial StrataMap technology.

## Ethics declarations

## Competing interests

N.F.F. holds stock options in Vizgen, Inc. A.B.S is employee of Center for Software Engineering Excellence (CSEE). The remaining authors declare no competing interests. This work was partially supported by a collaboration agreement between Illumina and the Broad Institute

## Supplementary Note

While Celldega was developed to visualize spatial transcriptomics data, its approaches are generalizable to additional data types. We demonstrate how we can use Celldega to visualize a single-cell gene expression dataset (45) obtained from the single-cell perturbation analysis library Pertpy (47) (Supp. Fig. 4A) We utilized a linked Clustergram and Enrich widget to visually explore data obtained from a CRISPRa activation experiment performed in a multipotent K562 cancer cell line (45). We observe that perturbations largely cluster according to their previously assigned biological program. Linked enrichment analysis enabled exploration of perturbation-associated gene sets; for example, as expected, genes upregulated after activation of erythroid differentiation-associated genes showed enrichment for erythroid and erythrocyte pathways (28) (Supp. Fig. 4A).

Celldega can also be extended beyond biological data, as demonstrated with public bicycle- share trip data from: Citi Bike NYC, Bluebikes Boston, Capital Bikeshare DC, and Divvy Chicago. We used Clustergram to identify highly connected neighborhoods from a station- by-station destination probability matrix and linked the results to a custom-built geospatial widget (Supp. Fig. 4B). This example illustrates Celldega’s generalizability while highlighting a shared challenge: entities can be similar in high-dimensional feature space while remaining constrained by and interpretable through their physical organization. Code and visualizations can be found here https://github.com/cornhundred/bike_network_traffic.

---

**Supp. Fig. 1.**
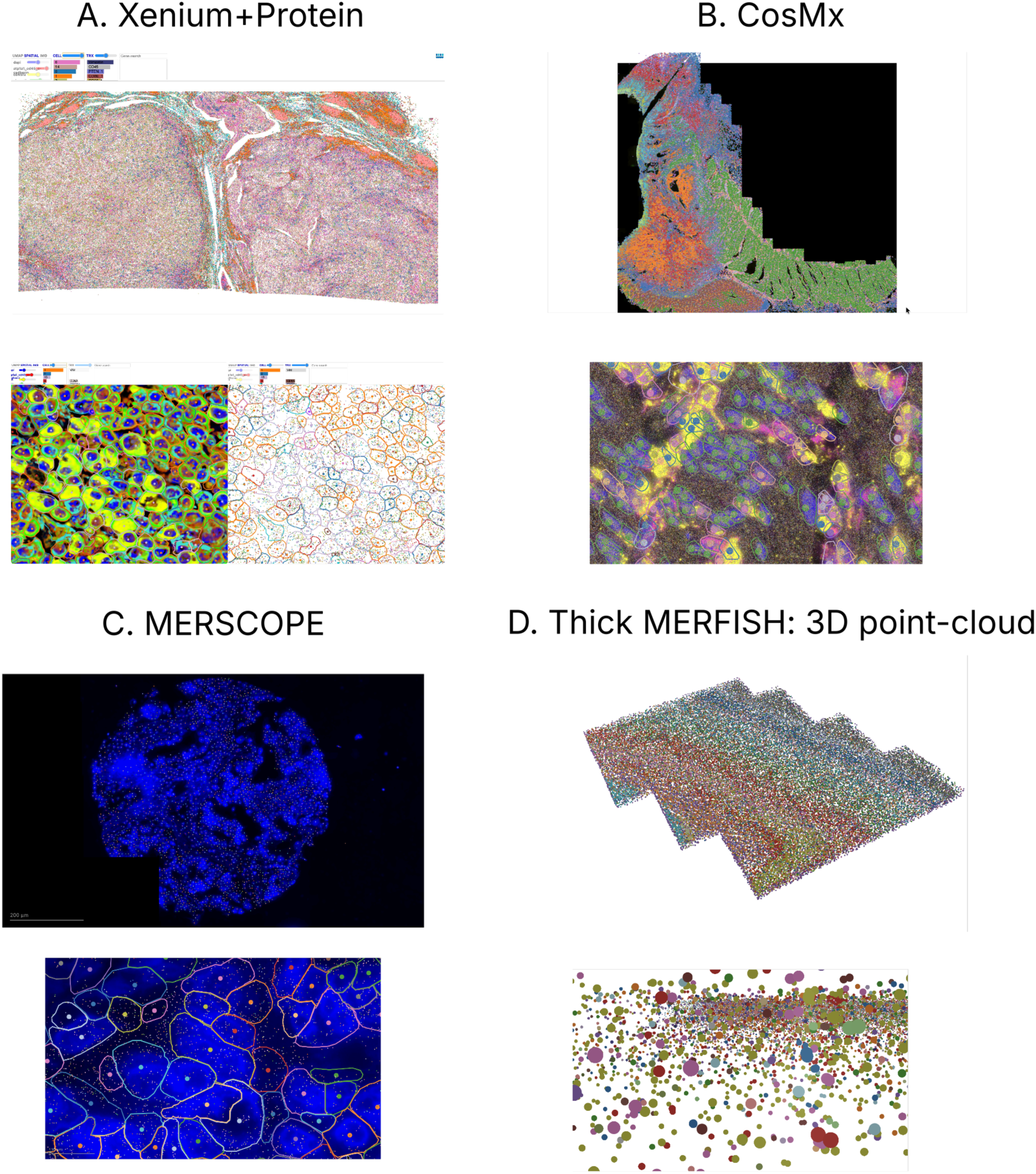
Additional Spatial Technologies. (A) Top, Landscape view of a publicly available human renal cell carcinoma Xenium+Protein dataset; bottom left, protein stain subcellular view; bottom right, transcript subcellular view. (B) Landscape view of a publicly available human colorectal cancer CosMx full-transcriptome dataset. (C) Landscape view of a tissue microarray (TMA) core of a human breast cancer MERSCOPE dataset. (D) Landscape 3D point-cloud visualization of a mouse brain thick MERFISH dataset.

**Supp. Fig. 2.**
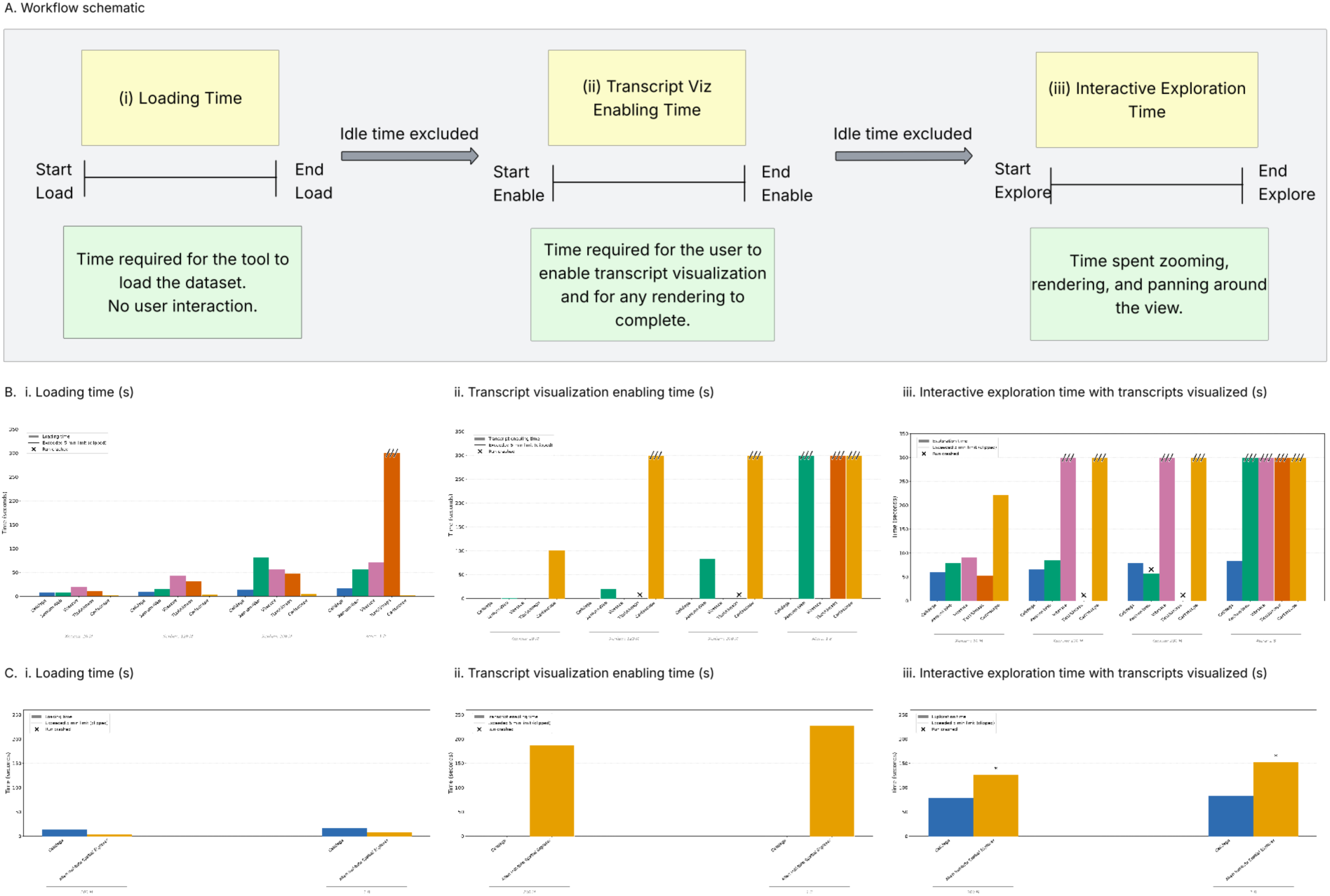
Celldega Performance Benchmark Against Spatial Visualization Tools. (A) Workflow schematic illustrating the three timed metrics. Loading time spans from the start to the end of the initial tool load, with no user interaction. Transcript enabling time spans from the start to the end of the transcript enabling step(s). Interactive exploration time covers the period during which the user zooms into a region, waits for all layers to render, and pans in four directions; this cycle was repeated across three regions per dataset. (B) Bar charts showing (i) average loading time, (ii) average transcript enabling time, and (iii) average interactive exploration time, each in seconds, across datasets of increasing size for Celldega, Cartoscope, Xenium Web, TissUUmaps, and Vitessce. Bars with a jagged top exceeded the 5-minute time limit and are clipped at 300 s. Black X markers indicate runs that crashed before completion. (C) Bar charts showing (i) average loading time, (ii) average transcript enabling time, and (iii) average interactive exploration time, each in seconds, for two datasets (200M and 1B transcripts) for Celldega and the Allen Institute Spatial Explorer. Celldega was run on Xenium and Atera datasets of roughly comparable scale to the two Atera datasets available through the Allen Institute Spatial Explorer. In panel (ii), for the Allen Institute Spatial Explorer, transcript enabling time reflects the time taken to manually enter 50 dataset-specific genes via copy-paste, which returned only a subset of the total transcripts present (approximately 5M and 20M transcripts for the 200M and 1B datasets, respectively). The * indicates that the Allen Instityte Spatial Explorer only visualized 50 genes rather than all genes (D) Formatted dataset size (in GB) for each tool— excluding the Allen Institute Spatial Explorer where we were unable to access the underlying data—across all four datasets.

**Supp. Fig. 3.**
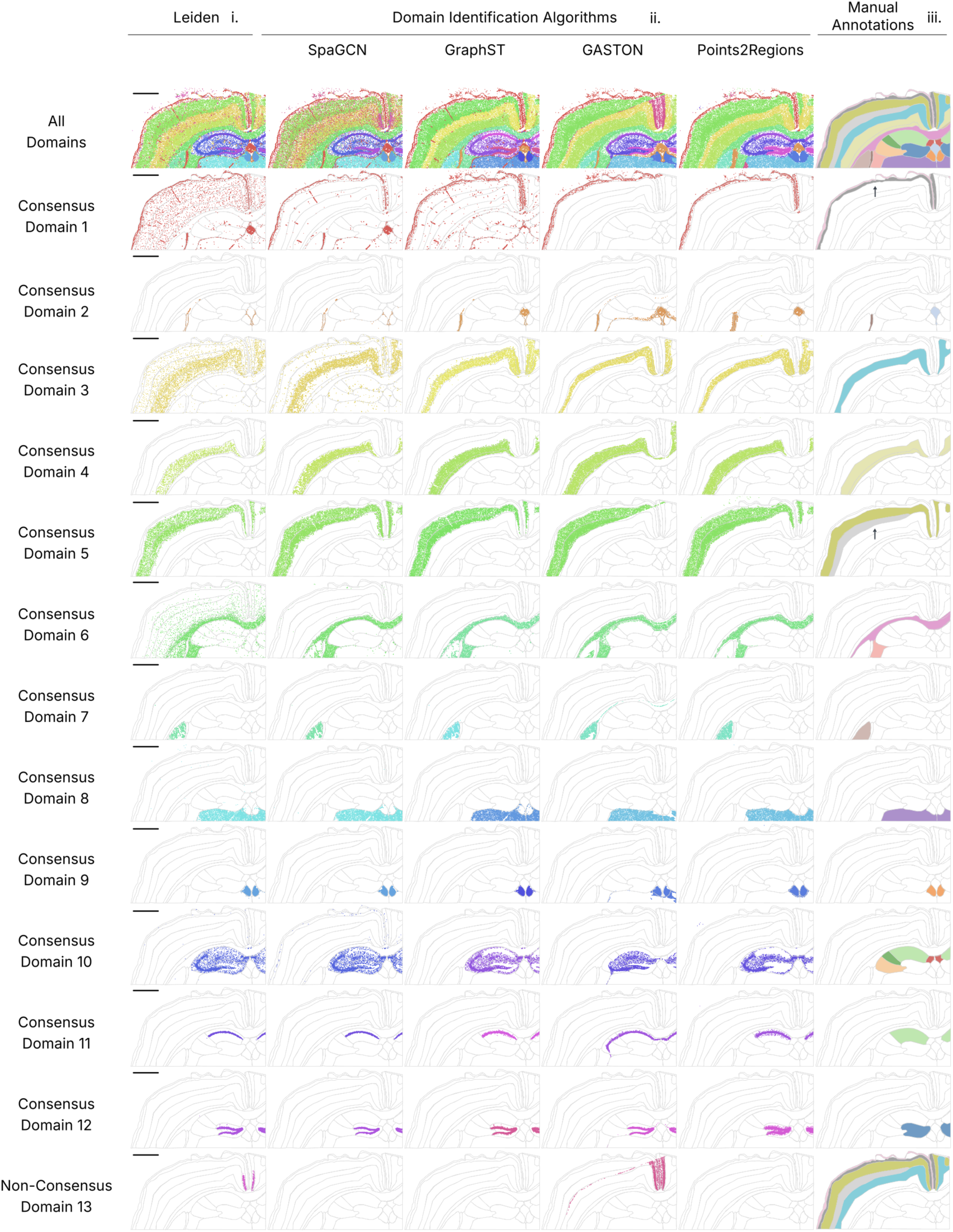
Consensus and Non-Consensus domain visualization across domain identification algorithms. Top row: Landscape visualization of Leiden clustering alongside domains identified across four independent algorithms (overlaid on manual annotation outlines) and manually annotated regions. Remaining rows: visualization of 12 commonly identified consensus domains and 1 unique non-consensus domain aligned with best fit manually annotated regions. The scale bars represent 1 mm. The arrows highlight (i) consensus domain 1 and consensus domain 5.

**Supp. Fig. 4.**
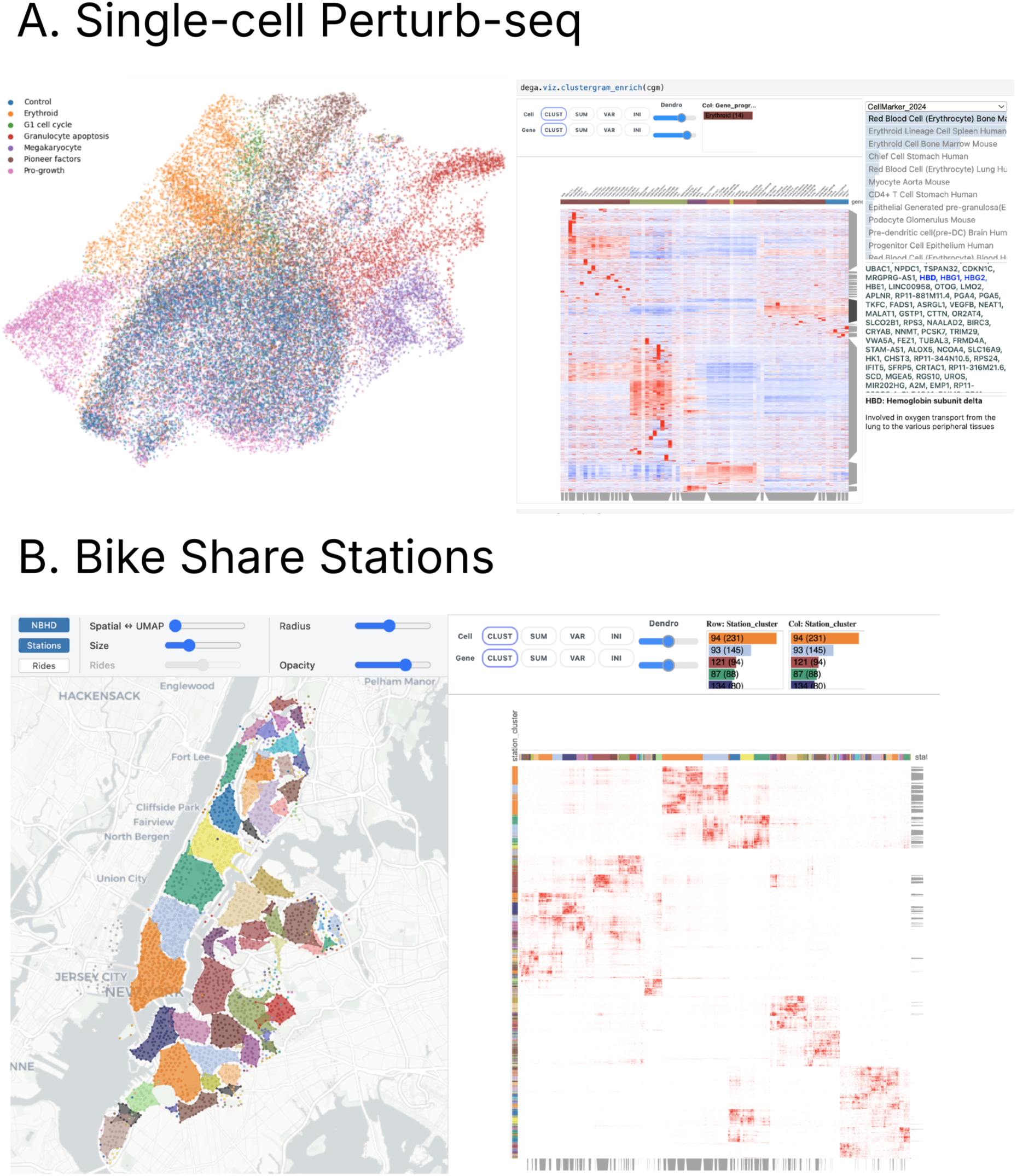
Non-spatial data. **(A)** UMAP, Clustergram, and Enrich visualization of CRISPRa perturb-seq single-cell gene expression signatures. Left, UMAP visualization of perturbation signatures colored by perturbation type. Middle, Clustergram visualization of hierarchical clustering of perturbation gene expression signatures. Right, Enrich enrichment analysis of erythrocyte associated gene module (CellMarker_2024) showing erythrocyte related gene set enrichment analysis results.

## Notes

https://broadinstitute.github.io/celldega

https://github.com/broadinstitute/celldega

### Summary of Updates

This version of the manuscript has been revised to remove a duplication of Fig. 2 .

